# Kinesin-8B controls basal body function and flagellum formation and is key to malaria parasite transmission

**DOI:** 10.1101/686568

**Authors:** Mohammad Zeeshan, David J. P. Ferguson, Steven Abel, Alana Burrrell, Edward Rea, Declan Brady, Emilie Daniel, Michael Delves, Sue Vaughan, Anthony A. Holder, Karine G. Le Roch, Carolyn A. Moores, Rita Tewari

**Author notes:** For correspondence Rita Tewari.

## Abstract

Eukaryotic flagella are conserved microtubule-based organelles that drive cell motility. *Plasmodium,* the causative agent of malaria, has a single flagellate stage: the male gamete in the mosquito. Three rounds of endomitotic division together with an unusual mode of flagellum assembly rapidly produce eight motile gametes. These processes are tightly coordinated but their regulation is poorly understood. To understand this important developmental stage, we studied the function and location of the microtubule-based motor kinesin-8B, using gene-targeting, electron microscopy and live cell imaging. Deletion of the *kinesin-8B* gene showed no effect on mitosis but disrupted 9+2 axoneme assembly and flagellum formation during male gamete development and also completely ablated parasite transmission. Live cell imaging showed that kinesin-8B-GFP did not colocalise with kinetochores in the nucleus but instead revealed dynamic, cytoplasmic localisation with the basal bodies and the assembling axoneme during flagellum formation. We thus uncovered an unexpected role for kinesin-8B in parasite flagellum formation that is vital for the parasite life cycle.

## Introduction

Eukaryotic flagella, also known as motile cilia, are conserved microtubule (MT)-based organelles that protrude from cells and drive motility of single cells, or the movement of fluid across ciliated tissue (Mirvis et al., 2018). In a number of organisms, the mechanisms by which flagella are built and maintained are gradually being revealed, and defects in flagellum assembly and function are associated with numerous human diseases (Baker and Beales, 2009; Croft et al., 2018)

In general, flagella grow from mother centrioles - also known as basal bodies - which are recruited from the centrosome to act as templates for the characteristic nine-fold symmetry of the MT array known as the flagellum axoneme (Mirvis et al., 2018). Specialised axonemal dynein motors are arranged according to the underlying axonemal MT organisation and these motors power flagella beating (Lin and Nicastro, 2018).Thus, flagella-driven motility depends on axonemal organisation defined by basal body function. However, the mechanisms that coordinate all these facets of flagella action are complex and less well understood.

Malaria is a disease caused by the unicellular parasite *Plasmodium* spp., which infects many vertebrates and is transmitted by female *Anopheles* mosquitoes (WHO, 2018). The parasite has a complex life cycle, with distinct asexual and sexual developmental stages. Extracellular, motile and invasive stages move by gliding motility powered by an actomyosin motor, except for male gametes which use flagellar movement (Boucher and Bosch, 2015; Frenal et al., 2017; Sinden et al., 1976; Sinden et al., 2010). Male gamete development from haploid gametocyte is a very rapid process with endomitotic division and flagellum formation completed in 10 to 15 minutes (Sinden et al., 2010). Specifically, three successive rounds of DNA replication produce an 8N nucleus and are accompanied by synthesis and assembly of basal bodies and axonemes; then cytokinesis and exflagellation release eight haploid flagellated male gametes (Sinden et al., 1976).

The *Plasmodium* male gamete has a very simple structure with no subcellular organelles except an axoneme with associated dyneins, elongated nucleus and surrounding flagellar membrane (Creasey et al., 1994; Okamoto et al., 2009; Sinden et al., 1976). The axoneme - essential for flagellar unit motility - consists of a pair of central MTs (C1 and C2) encircled by nine doublet MTs – a so-called 9+2 organisation - as in many other organisms (Mitchell, 2004). However, the *Plasmodium* flagellum differs in several key respects in the mechanism of axoneme formation from that of other organisms including trypanosomes*, Chlamydomonas* and humans (Bastin, 2010; Briggs et al., 2004; Soares et al., 2019), as well as other Apicomplexa with flagellated gametes like *Toxoplasma* and *Eimeria* (Ferguson, 2009; Ferguson et al., 1977; Ferguson et al., 1974). The differences in the mechanism of axoneme formation are: 1) the *Plasmodium* basal body, from which the flagellum is assembled, does not exhibit the classical nine triplet MT (9+0) (Mirvis et al., 2018), but instead consists of an electron dense amorphous structure within which nine peripherally arranged MTs can sometimes be resolved (Francia et al., 2015); 2) basal bodies/centrioles are not present during asexual stages of the *Plasmodium* life cycle, unlike Toxoplasma and Eimeria where they play a role during asexual division (Ferguson DJP, 2013), but assemble de novo in male gametocytes (Sinden et al., 1976); 3) the axoneme itself is assembled from the basal bodies within the male gametocyte cytoplasm, and only after its assembly does the axoneme protrude from the membrane to form of the flagellum itself; as a result and very unusually, axoneme construction occurs independently of intra-flagellar transport (IFT)(Briggs et al., 2004; Sinden et al., 1976). Mechanisms that regulate *Plasmodium* male gamete biogenesis and control axoneme length are largely unknown, although the involvement of some individual proteins has been described. For example, PF16 is a flagellar protein with an essential role in flagellar motility and the stability of the central MT pair in *Chlamydomonas* and *Plasmodium* (Smith and Lefebvre, 1996; Straschil et al., 2010), while SAS6, a well-conserved basal body protein, has a role in *Plasmodium* flagellum assembly (Marques et al., 2015; Nigg and Stearns, 2011). Proteomic analysis has revealed many potential regulators of axoneme assembly present in male gametes, including members of the kinesin superfamily (Talman et al., 2014). Kinesins are molecular motor proteins that have essential roles in intracellular transport, cell division and motility (Cross and McAinsh, 2014; Verhey and Hammond, 2009; Wittmann et al., 2001). Some kinesin families are known to contribute to ciliary structure and function, either by transporting ciliary components along axonemal MTs by IFT like kinesin-2s (Scholey, 2008), or by destabilizing MTs, like kinesin-9 and kinesin-13 (Blaineau et al., 2007; Dawson et al., 2007).

Kinesin-8s are conserved from protozoa to mammals, and can be sub-classified as kinesin-8A, −8B and −8X (Vicente and Wordeman, 2015; Wickstead et al., 2010). In general, kinesin-8s are multi-tasking motors, able to move along MTs, cross-link and slide MTs and influence MT dynamics at their ends (Mayr et al., 2007; Su et al., 2013). Kinesin-8As are best characterised as regulators of spindle length and chromosome positioning during metaphase (Mary et al., 2015; Savoian et al., 2004; Straight et al., 1998), whilst the mammalian kinesin-8B Kif19has a role in ciliary length control (Niwa et al., 2012).

The *Plasmodium* genome encodes two kinesin-8s (kinesin-8X and kinesin-8B), along with 6 other kinesin proteins (Vicente and Wordeman, 2015; Wickstead et al., 2010; Zeeshan M, 2019). Kinesin-8X is conserved across the Apicomplexa whereas kinesin-8B is restricted to certain genera such as *Toxoplasma*, *Eimeria* and *Plasmodium* that have flagellated gametes (Zeeshan M, 2019). However, the role of kinesin-8s in regulation of *Plasmodium* flagella is unknown.

Here we have analysed the location and function of the kinesin-8B protein (PBANKA_0202700) present in male gametes in the rodent malaria parasite, *Plasmodium berghei*. Deletion of the gene results in complete impairment of 9+2 axoneme formation because of misregulation of the basal body, and no effect on nuclear division. Its unprecedented involvement in basal body function prevents assembly of axonemes and the motile flagellum, and thereby blocks parasite transmission. Live cell imaging showed that kinesin-8B-GFP is expressed only during male gamete development and that strikingly, it is not associated with the nuclear kinetochore but is present only in the cytoplasm. Here, kinesin-8B-GFP dynamically associated with the basal bodes and with axonemes during their formation and assembly, consistent with an unexpected role in flagellum genesis.

## Results

### Kinesin-8B is essential for male gamete formation and its deletion blocks parasite transmission

Based on its known expression in male gametes and a potential function during male gamete development, we first assessed the function of kinesin-8B throughout the *Plasmodium* life cycle by using a double crossover homologous recombination strategy to delete its gene (S1A Fig). Diagnostic PCR confirmed successful integration of the targeting construct at the *kinesin-8B* locus (S1B Fig), and analysis by qRT-PCR confirmed complete deletion of the *kinesin-8B* gene in this transgenic parasite (S1C Fig). Successful deletion of the *kinesin-8B* gene indicated that it is not essential for asexual blood stage development, consistent with the protein’s expression and localisation only during male gametogenesis, and with global functional studies (Bushell et al., 2017). Phenotypic analysis of the *Δkinesin-8B* parasite was carried out at other developmental stages, comparing the wild type (WT-GFP) parasite with two independent gene-knockout parasite clones generated by two different transfections. Since *Δkinesin-8B* parasites undergo asexual blood stage development, form gametocytes in mice, and exhibit no change in morphology or parasitemia, an essential role for kinesin-8B during these stages is unlikely. We then examined male and female gametocytes following activation in exflagellation/ookinete medium for the development and emergence of male and female gametes. This analysis revealed that there was no exflagellation in either of the *Δkinesin-8B* parasite clones (Clone 1 and clone 4), whereas exflagellation frequency for the WT-GFP parasites was normal (Figure 1A).

**Figure 1.**
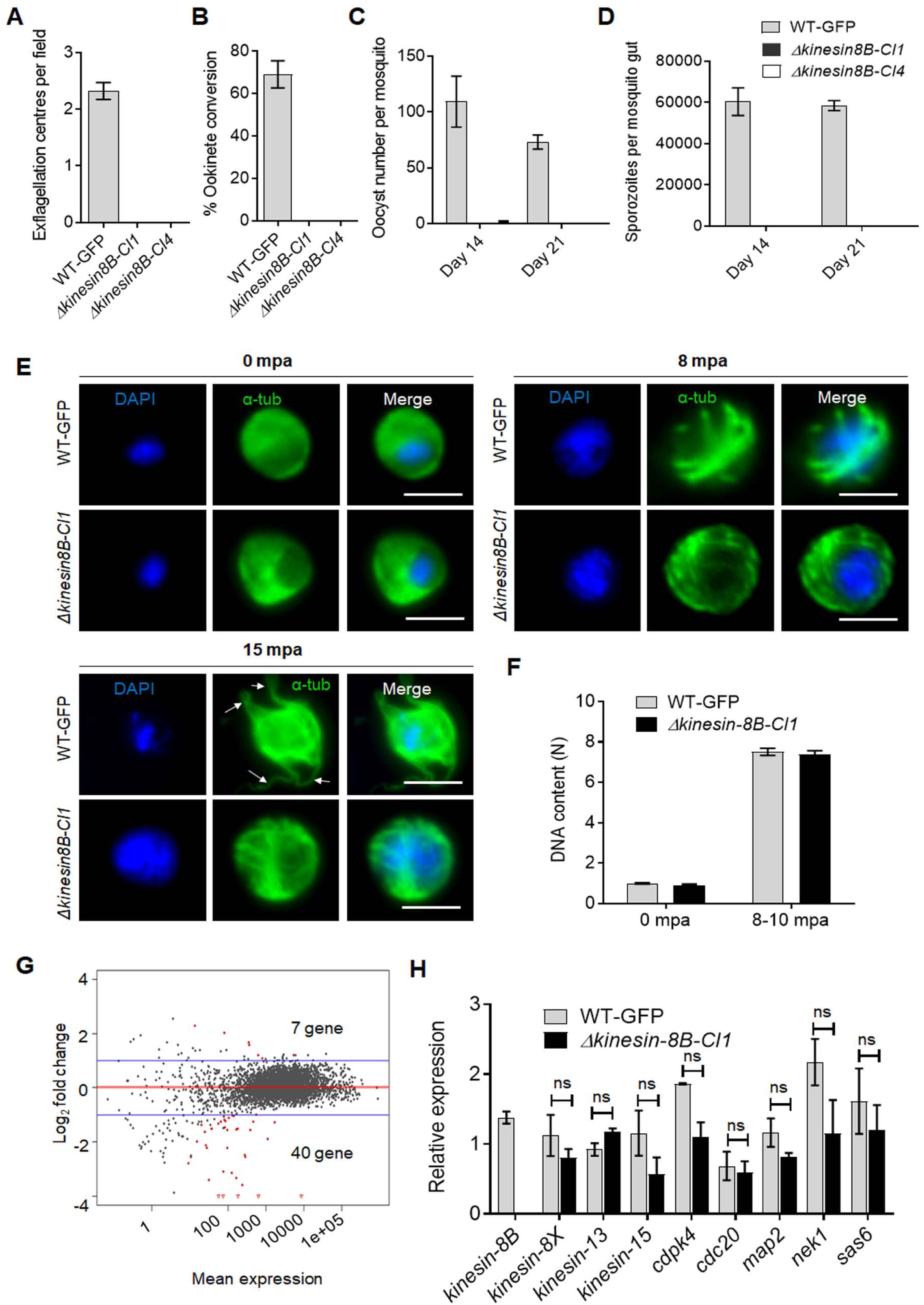
Kinesin-8B is required for male gamete formation and parasite transmission. **A**. Male gametes (determined by quantifying exflagellation centres) are absent in Δ*kinesin-8B* clones (Cl1 and Cl4) but not wild type parasites. Data are means from 15 random independent fields with a 40x objective, and three independent replicates. Error bars: ± SEM. **B**. Ookinete conversion from zygotes in wildtype and *Δkinesin-8B* parasites (Cl1 and Cl4). n = >3 independent experiments (>70 cells per experiment for wild type). Error bars: ± SEM. **C**. Total number of oocysts of *Δkinesin-8B* (Cl1 and Cl4) and WT-GFP parasites in mosquito midguts at 14 and 21 dpi. Data are shown as Mean ± SEM, n = 3 independent experiments. **D**. Total number of sporozoites in oocysts of *Δkinesin-8B* (Cl1 and Cl4) and WT-GFP parasites at 14 and 21 dpi. Mean ± SEM. n = 3 independent experiments. **E**. In the absence of kinesin-8B, α-tubulin is not reorganised during gametogenesis. IFA images of α-tubulin in WT-GFP and *Δkinesin-8B* male gametocytes either before activation (0 mpa) or 8 and 15 mpa. DAPI (DNA) is blue Scale bar = 5 µm. **F**. Fluorometric analyses of DNA content (N) after DAPI nuclear staining. Male gametocytes were at 0 min (non-activated), or 8-10 min post activation. The mean DNA content (and SEM) of 10 nuclei per sample are shown. Values are expressed relative to the average fluorescence intensity of 10 haploid ring-stage parasites from the same slide **G**. Differentially expressed genes in *Δkinesin-8B* parasites compared to WT-GFP parasites. Samples were collected at 0 mpa. **H**. qRT-PCR analysis of changes in transcription of selected genes affected in *Δkinesin-8B* parasites. Error bars: ± SEM. The selected genes have established or probable role in male gamete development. ns: non-significant.

To investigate this defect further, we examined both zygote formation and ookinete development. No zygote (or ookinete) formation was detected in either clonal *Δkinesin-8B* parasite line, whereas WT-GFP parasites showed normal zygote (ookinete) development (Figure 1B). To examine the effect of this defect on parasite transmission, *Anopheles stephensi* mosquitoes were fed on mice infected with *Δkinesin-8B* parasites, and the number of GFP-positive oocysts on the mosquito gut was counted 14 days later. No oocysts were detected in mosquito guts infected with *Δkinesin-8B* parasites, except on one occasion when we found 4 small oocysts in one mosquito after 14 days of infection with the clone 4 parasite line (Figure 1C). After 21 days of infection, no oocysts were observed for the *Δkinesin-8B* lines whereas the WT-GFP parasites line produced normal growing oocysts (Figure 1C). No sporozoites were found in the oocysts of mosquitoes fed with *Δkinesin-8B* parasites neither at 14 days post infection (dpi) nor 21 dpi (Figure 1D).

### Kinesin-8B is not required for DNA replication but its deletion impairs male exflagellation

To understand the defect in more detail, we analysed the development of MT structures in *Δkinesin-8B* and WT-GFP male gametocytes following activation *in vitro* by treatment with xanthurenic acid and decreased temperature at 0, 8 or 15 minutes post activation (mpa). We observed no very clear early differences, but the MT distribution in the *Δkinesin-8B* gametocytes gradually diverged from that of the WT-GFP parasites over time, and these mutants did not form flagella (as evidenced from tubulin labelling) at 15 mpa **(arrow**, Figure 1E). To assess the effect of the mutations on DNA replication during male gametogenesis, we analysed the DNA content (N) of *Δkinesin-8B* and WT-GFP male gametocytes by fluorometric analyses after DAPI staining. We observed that *Δkinesin-8B* male gametocytes had a haploid DNA content (1N) at 0 min (non-activated) and were octaploid (8N) 8-10 min post activation, similar to WT-GFP, indicating the absence of kinesin-8B had no effect on DNA replication (Figure 1F).

### Transcriptomic analysis identified no additional genes responsible for the Δ*kinesin-8B* phenotype

To investigate further the defect in male gamete development in *Δkinesin-8B* parasites, we analysed the transcript profile of both non-activated and activated *Δkinesin-8B* and control WT-GFP gametocytes. RNAseq analysis was performed on WT-GFP and *Δkinesin-8B* gametocytes, just before activation (0 mpa) and after exflagellation (30 mpa)) for four pairs of biological replicates (WT, 0 mpa; WT, 30 mpa; *Δkinesin-8B,* 0 mpa; and *Δkinesin-8B,* 30 mpa) and the read coverages exhibited Spearman correlation coefficients of 0.97, 0.98, 0.96, and 1.00, respectively, demonstrating the reproducibility of this experiment. As expected, no significant reads mapped to the *Δkinesin-8B* region (S1D Fig). Only 7 genes were upregulated, and 40 genes were downregulated compared to the WT-GFP control, but none was known to be important in gametogenesis (Figure 1G, **S1 Table)**. In a qRT-PCR analysis of a specific set of genes coding for proteins identified either as molecular motors (kinesin family) or involved in male gamete development (Billker et al., 2004; Ferguson DJP, 2013; Tewari et al., 2005) (Figure 1H) showed no significant change in transcript level. These data suggest that the observed phenotype resulted directly from the absence of kinesin-8B rather than through an indirect effect on another gene transcript.

### Activated *Δkinesin-8B* gametocytes lack the 9+2 MT architecture of axoneme

To examine ultrastructural differences between the *Δkinesin-8B* and WT-GFP parasite lines during male gametogenesis, cells at 8 and 15 mpa were examined by electron microscopy. The early developmental stage of both parasite lines contained a large central nucleus with diffuse chromatins (Figure 2.1a, b). At 8 mpa, elongated axonemes were identified running round the peripheral cytoplasm of the WT-GFP parasite (Figure 2.1a, c), and in contrast, the cytoplasm of the mutant contained numerous elongated but randomly oriented MTs (Figure 2.1b, d). The nuclear appearance of mutant and WT-GFP parasites was similar in mid-stage male gamete development (Figure 2.1e, f); both had similar electron dense cone-shaped nuclear poles from which MTs radiated to form the nuclear spindle with attached kinetochores (Figure 2.1g, h), the association between the basal bodies and nuclear poles were dramatically reduced in *Δkinesin-8B* mutant **(cf**. Figure 2.1g, h).

In WT-GFP parasites several axonemes with the classical 9+2 arrangement of duplet and single MTs were observed in the cytoplasm (Figure 2.2a **and insert)**, although some abnormal and incomplete axonemes were also observed (Figure 2.2a, **arrows)**. In marked contrast, the cytoplasm of the mutant parasite contained numerous randomly distributed duplet and single MTs with no overall structural organization (Figure 2.2b **and Insert)**. In the late stages of development (15 mpa), it was possible to observe chromatin condensation in the nuclei of WT-GFP and *Δkinesin-8B* parasites, consistent with successful endomitosis in the mutant (Figure 2.2c, d). In the WT-GFP parasite at 15 mpa, male gamete formation and exflagellation was observed, with the protrusion from the surface of the male gametocyte of a flagellum together with its attached nucleus leaving a residual body of cytoplasm (Figure 2.2c, e). Free male gametes consisting of an electron dense nucleus and associated 9+2 flagellum were also observed (Figure 2.2e **insert)**. In contrast, in the *Δkinesin-8B* mutants at 15 mpa, there was little change in the cytoplasm, which still contained large numbers of randomly distributed MTs (Figure 2.2f, **insert)** and no cytoplasmic processes or male gamete formation were observed.

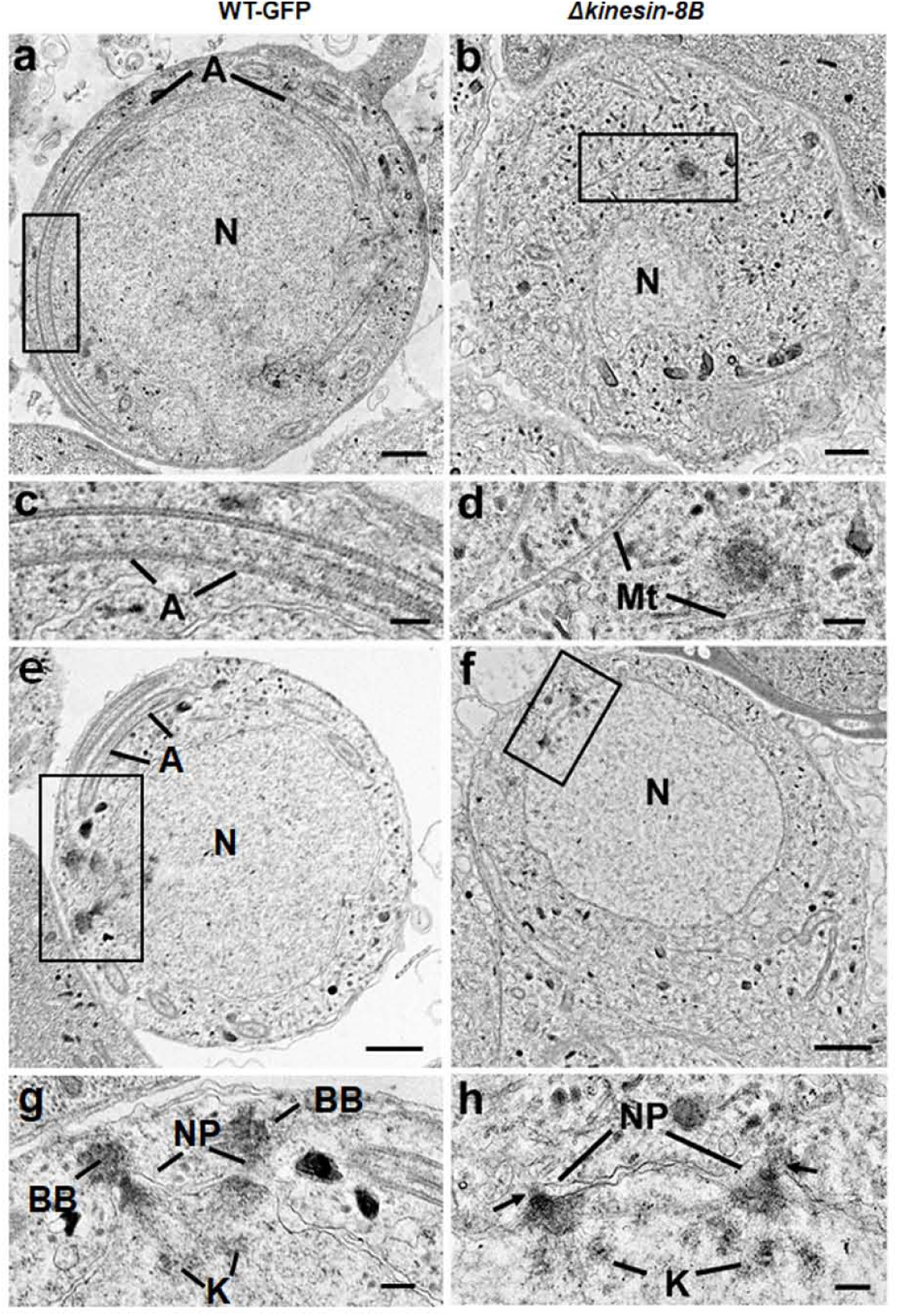
2. Ultrastructure analysis reveals defects in basal body and axoneme formation: **2.1**. Electron micrographs of mid-stage (8 mpa-min post activation) male gametocytes of WT-GFP (**a,c,e,g**) and *Δkinesin-8B* (**b,d,f.h**) parasites. Bars represent 1µm in **a, b, e** and **d** and 100 nm in **c, d, g** and **h**. **a**. Low power image of a WT-GFP male gametocyte showing the large nucleus (N) with cytoplasm containing long axonemes (A) running around the periphery. **b**. Low power image of a *Δkinesin-8B* male gametocyte showing the nucleus (N) and the cytoplasm containing randomly orientated microtubules (Mt). **c**. Detail of the enclosed area in panel **a** showing the parallel organization of the microtubules forming an axoneme (A). **d**. Detail of the enclosed area in panel **b** showing randomly orientated microtubules (Mt). **e**. WT-GFP male gametocyte showing the large nucleus (N) with multiple nuclear poles, spindle and basal bodies with cytoplasm containing a number of axonemes (A). **f**. *Δkinesin-8B* male gametocyte showing the nucleus with nuclear poles and spindle but no basal bodies. The cytoplasm lacks axonemes but has numerous microtubules (Mt). **g**. Detail from the enclosed are in **e** showing basal bodies (BB), the nuclear pole (NP) connected by spindle microtubules with attached kinetochores (K). **h**. Detail from the enclosed area in **f** showing the nuclear poles (NP) connected by spindle microtubules with attached kinetochores (K). Note the absence of basal bodies associated with the nuclear poles (arrows).

**2.2.**
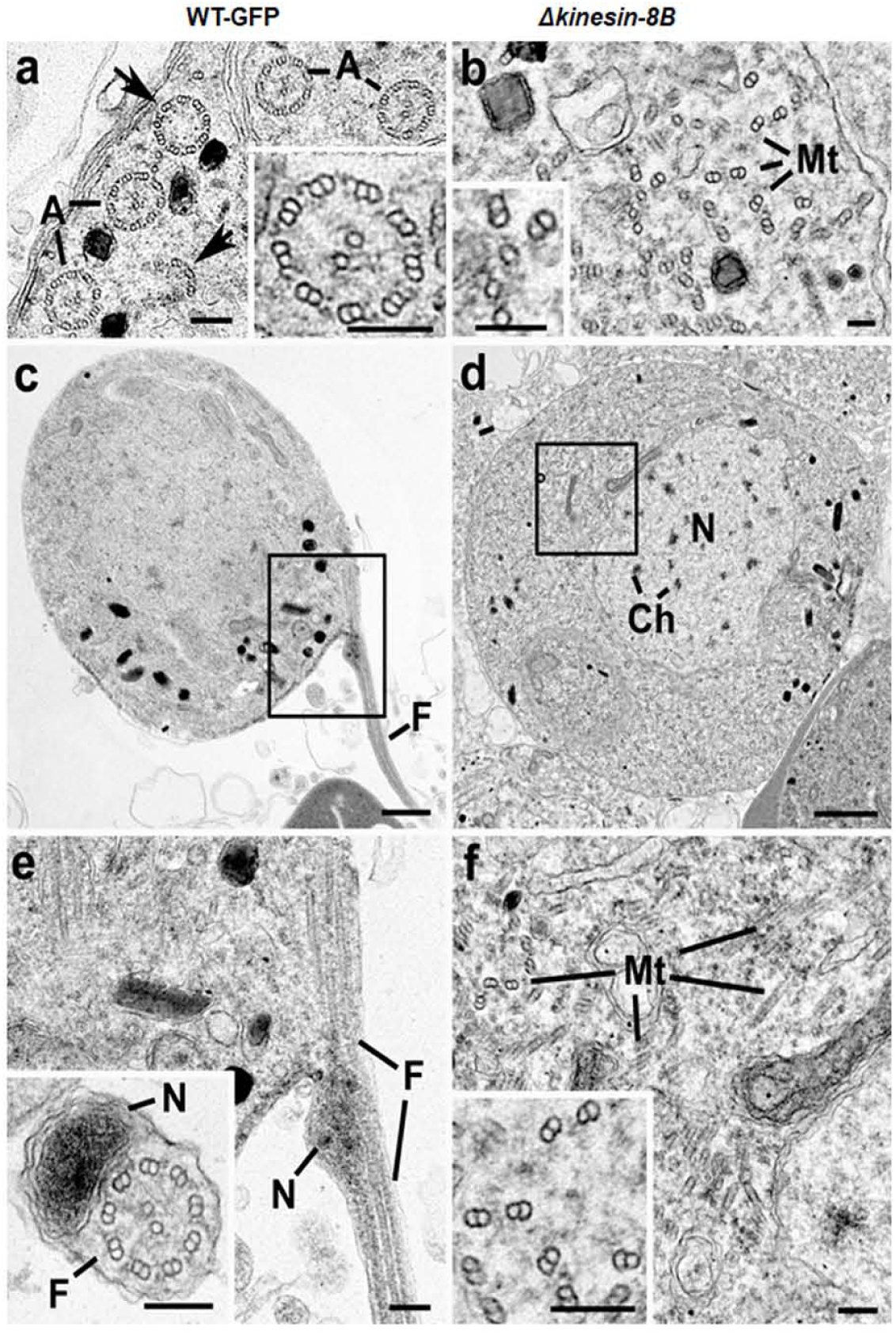
Electron micrographs of mid (8 mpa) and late (15 mpa) male gametocytes of WT-GFP (**a,c.e**) and mutant (**b,d,f**). Bar represent 1 µm in **c** and **d** and 100 nm in all other micrographs. **a**. Part of the peripheral cytoplasm of a mid-stage (8 mpa) WT-GFP gametocyte showing a cross section through a number 9+2 axonemes (A). Note the presence of incomplete axonemes (arrows). **Insert**. Detail of a cross section axoneme showing the 9+2 organisation of the microtubules. **b**. Part of the cytoplasm of a mid-stage mutant male gametocyte showing numerous randomly orientated microtubules (Mt). **Insert**. Detail showing the presence of both duplet and single microtubules. **c**. Late stage (15 mpa) male gametocyte showing the partial formation of a male gamete by exflagellation. F – Flagellum. **d**. Late mutant male gametocyte showing the early chromatin condensation and the absence of axonemes in the cytoplasm. **e**. Detail of the enclose area in **c** showing the flagellum (F) and associated nucleus (N) protruding from the surface of the male gametocyte. **Insert**. Cross-section through a free male gamete showing the electron dense nucleus (N) and the classical 9+2 flagellum. **f**. Detail of the enclosed in **d** showing the numerous randomly orientated microtubules (Mt). **Insert**. Detail showing the disorganisation of the duplet microtubules.

In summary, deletion of the *kinesin-8B* gene did not appear to affect the initiation or growth of single and duplet MTs but association of basal bodies with the nuclear pole appeared to be reduced and coordinated formation of the 9+2 organisation of functional axonemes did not occur. This defect prevented exflagellation and the formation of viable male gametes.

### Spatio-temporal profiles of kinesin-8B and the kinetochore protein Ndc80 show that kinesin-8B is absent from the mitotic spindle

To understand more precisely the role of kinesin-8B in building the male gametocyte flagellum, we investigated the subcellular location of kinesin-8B by live cell imaging in *P. berghei*. We generated a transgenic parasite line by single crossover recombination at the 3’ end of the endogenous *kinesin-8B* locus to express a C-terminal GFP-tagged fusion protein (S2A Fig). PCR analysis of genomic DNA using locus-specific diagnostic primers indicated correct integration of the GFP tagging construct (S2B Fig), and the presence of a protein of the expected size (∼198 kDa) in a gametocyte lysate was confirmed by western blot analysis using GFP-specific antibody (S2C Fig). The expression and location of kinesin-8B was assessed by live cell imaging throughout the parasite life cycle, and it was observed only in male gametocytes and gametes.

In order to establish whether kinesin-8B is part of the mitotic assembly during the three rounds of chromosome replication in male gamete development, we examined its location together with that of the kinetochore protein Ndc80. Parasite lines expressing kinesin-8B-GFP and Ndc80-cherry were crossed and used for live cell imaging of both markers to establish their spatio-temporal relationship. 1-2 min after gametocyte activation, kinesin-8B was observed close to the nucleus and adjacent to Ndc80, but not overlapping with it (Figure 3A). This pattern of adjacent location is also seen in later stages of development. This shows that both axoneme formation and chromosome division begin at very early stage of gametogenesis and continue side by side (Figure 3A). Completion of both processes is coordinated in a spatio-temporal manner before the onset of exflagellation, but kinesin-8B’s role is specifically related to axoneme formation in the cytoplasm.

**Figure 3.**
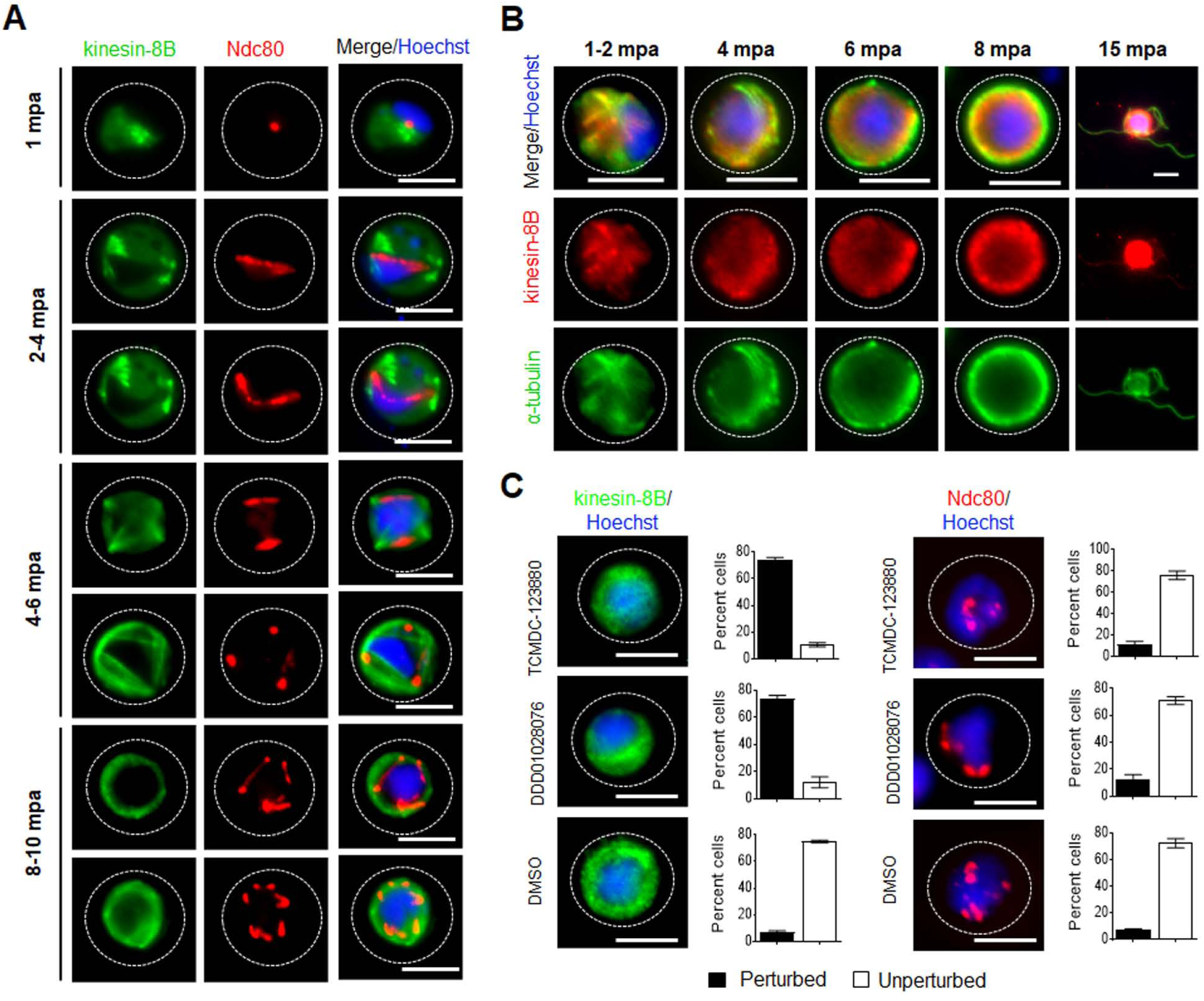
The location of kinesin-8B in relation to that of the kinetochore (Ndc80) and microtubule (α-tubulin) markers. **A**. The location of kinesin-8B-GFP (green) in relation to the kinetochore marker, Ndc80-mCherry (red) during male gamete formation. The cytoplasmic location of kinesin-8B contrasts with the nuclear location of Ndc80 during chromosome replication and segregation, indicating that kinesin-8B is not associated with the mitotic spindle. **B**. Indirect immunofluorescence assay (IFA) showing co-localization of kinesin-8B (red) and α-tubulin (green) in male gametocytes at 1 to 2, 4, 6, 8 and 15 mpa. **C**. Anti-malarial molecules block the dynamic distribution of kinesin-8B showing the resulting phenotype of compound addition at 4 mpa. In contrast to the effect on kinesin-8B-GFP distribution no significant effect was seen on Ndc80–RFP. Inhibitors were added at 4 mpa and parasites were fixed at 8 mpa. Scale bar = 5 µm.

### Kinesin-8B localizes to cytoplasmic MTs and its movement is blocked by antimalarial compounds

To further explore the location of kinesin-8B, we investigated its co-localization with MTs (using α-tubulin as a marker) by indirect immunofluorescence assay (IFA) using gametocytes fixed at different time points after activation. Kinesin-8B was localized on cytoplasmic MTs rather than on spindle MTs during male gamete formation. The pattern of both MTs and kinesin-8B was more distinct in later stages where their distribution adopted an axoneme-like pattern around the nucleus (Figure 3B). Further we tested two molecules with known antimalarial properties (TCMDC-123880 and DDD01028076) with a putative role in targeting MT dynamics during male gamete development (Delves et al., 2018; Gamo et al., 2010) Addition of these molecules at 4 mpa blocked the dynamic distribution of kinesin-8B in over 80% of male gametocytes had no significant effect on Ndc80 dynamics (Figure 3C). This suggest that these compounds are most effective against cytoplasmic MT, and serve to distinguish the cytoplasmic and nuclear MT-based processes.

### Kinesin-8B associates dynamically with basal bodies and growing axonemes

Since the *Δkinesin-8B* parasite appeared to have defects in basal body interactions with nuclear pole and axoneme assembly, we analysed further the dynamic location of kinesin-8B by live imaging. We observed a diffuse cytoplasmic localization in non-activated gametocytes, and following activation kinesin-8B accumulated at one side of the nucleus, while retaining a cytoplasmic location. Within 1 to 2 minutes after activation, the distribution of kinesin-8B showed four clear foci close to the nucleus, suggesting its association with the ‘tetrad of basal bodies’ which is the template for axoneme polymerization (Figure 4A) (Sinden et al., 1978; Sinden et al., 1976). These four foci duplicated further within 2-4 min to form another set of four kinesin-8B foci, which later moved apart from each other (Figure 4A). As gametogenesis proceeds, these kinesin-8B foci continued to move apart forming fibre-like structures around the nucleus by 4-6 mpa. The growth of these kinesin-8B-tagged fibres resembles the growing cytoplasmic axonemes (Marques et al., 2015; Straschil et al., 2010). The duplication of the four foci and emergence of these fibre-like structures occurred within 2-4 min (Figure 4B, **Supplementary videos SV1 and SV2)**. By 6-8 mpa, these fibre-like structures decorated by kinesin-8B were arranged in a specific pattern with the basal bodies around the nucleus and forming a basket-like structure, similar to that described in earlier ultrastructural studies (Sinden et al., 1978; Sinden et al., 1976). This structure likely corresponds to completed axonemes. Kinesin-8B was also observed with a continuous distribution along the length of the gametes as they emerged from the residual body of the gametocyte. Overall, the spatial and temporal profile of kinesin-8B revealed a localisation to basal bodies and axonemes throughout gametogenesis.

**Figure 4.**
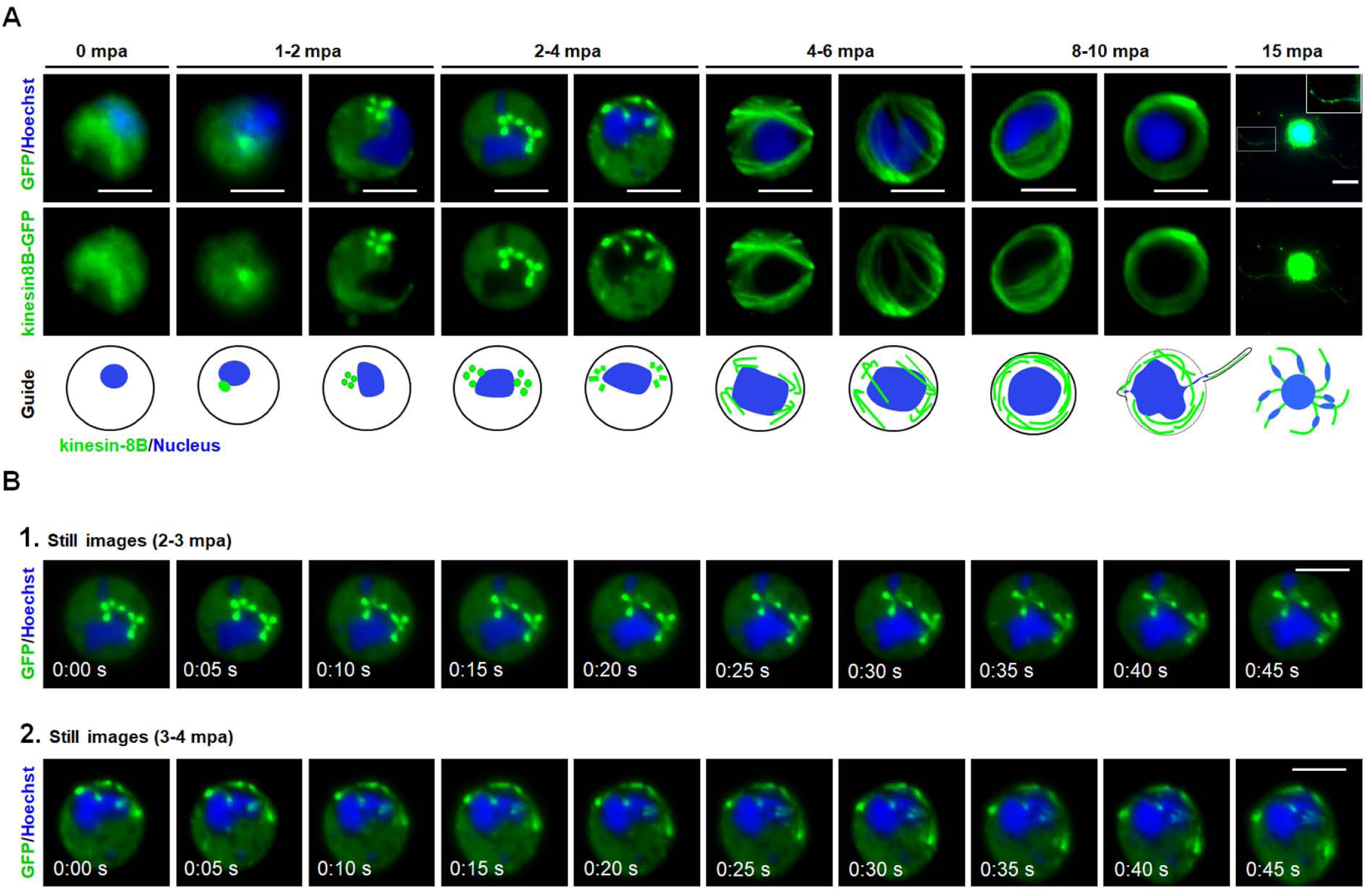
The dynamic location of kinesin-8B in basal body and axoneme formation. **(A)** Cytoplasmic location of kinesin-8B-GFP (green) by live cell imaging in real time during male gamete development. DNA is stained with Hoechst 33342 (blue). Before activation (0 min) kinesin-8B has a diffused location in male gametocyte cytoplasm. One to two min post activation (mpa) it accumulates at one end of the nucleus and forms four foci reminiscent of the ‘tetrad of basal body’ defined by electron microscopy (Sinden et al, 1976). These tetrad foci are duplicated within 2 to 4 mpa, and by 4 to 6 mpa fibre-like structures, representing axonemal growth decorated with kinesin-8B-GFP, extend from these tetrads to make a basket-like structure around the nucleus, which is completed by 8 to 10 mpa. Following exflagellation kinesin-8B-GFP is located along the length of the flagellum in the free male gamete (15 mpa, inset). For each time point a cartoon guide is presented. **B**. Still images (at every 5 seconds) of tetrad-foci duplication and the start of axonemal growth decorated with kinesin-8B-GFP at 2 to 3 mpa (Fig 3B.1; Supplementary videos SV1) and 3 to 4 mpa (Fig 3B.2; SV2). Scale bar = 5 µm.

Based on the ultrastructural studies, live cell imaging, and immunofluorescence microscopy, we suggest a model for the dynamic location of kinesin-8B on basal bodies and axonemes during male gamete development, which is depicted in Figure 5.

**Figure 5.**
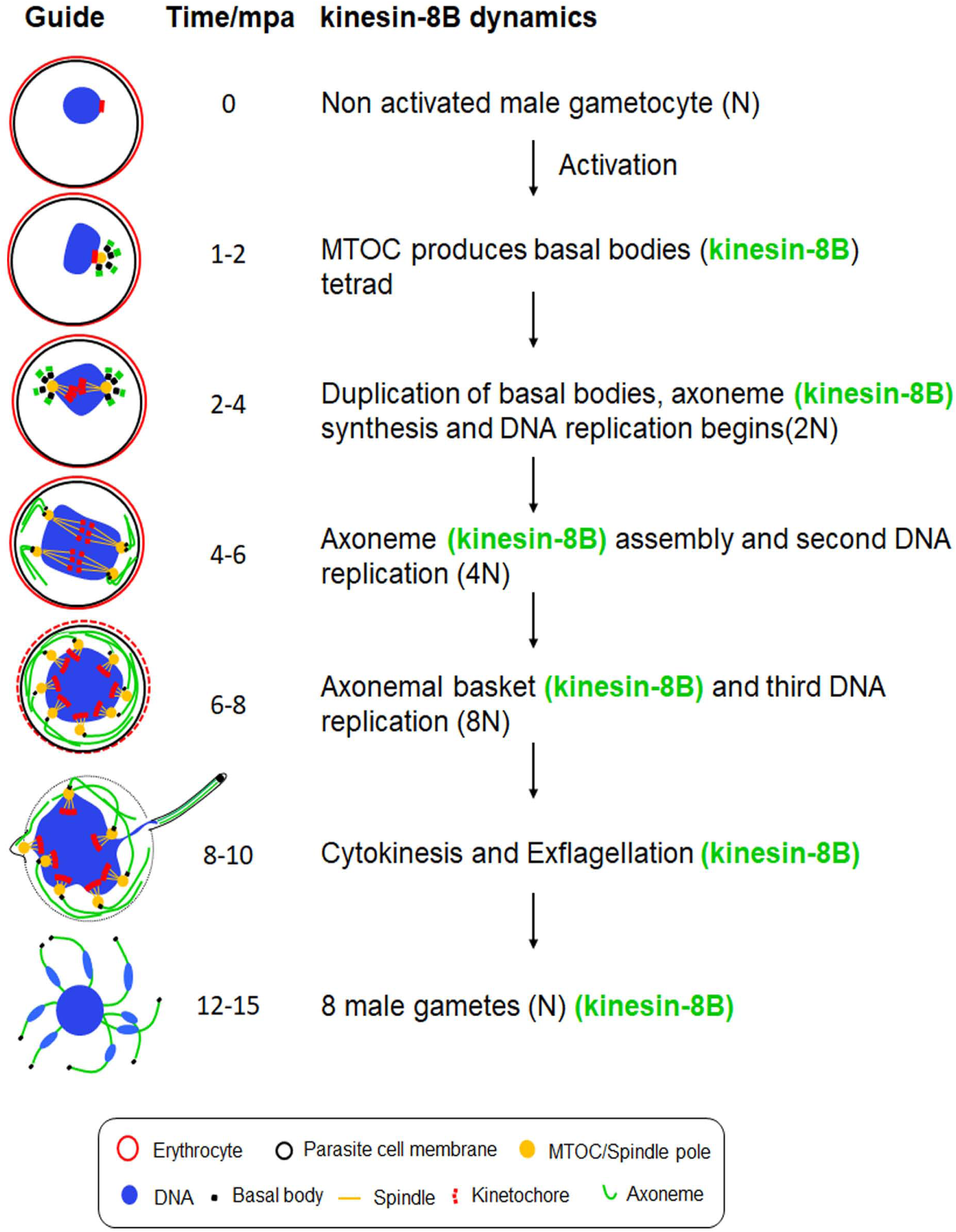
Schematic model of kinesin-8B location on basal bodies and axonemes assembly during the male gamete development. Kinesin-8B accumulates at one end of nucleus within 1 min post activation (mpa) of male gameyocytes, probably near the MT organizing centre (MTOC), and develops into four foci corresponding to the tetrad of basal bodies (BBs, green). These tetrad foci of kinesin-8B duplicate and separate from each other within 2-4 mpa. Each of these foci is a basal body that serves as a platform for axonemal MT assembly. The fibre-like structure decorated with kinesin-8B (green) emerges from these foci showing their association with axoneme assembly. Meanwhile, DNA replication accompanies basal body duplication, with associated mitotic spindles (yellow) and kinetochores (red) via the spindle pole (Sinden et al., 1976). The axonemal fibre-like structures develop further into a basket-like structure around the nucleus by 6-10 mpa. Axoneme formation and DNA replication are completed by this time (8-10 mpa) and a haploid genome connected with a basal body, is pulled into an emerging gamete.

## Discussion

Flagella and cilia, built around a central array of axonemal MT, are ancient organelles involved in motility and signalling (Carvalho-Santos et al., 2011; Soares et al., 2019). In some Apicomplexa parasites, including *Plasmodium* only the male gamete is flagellate in the sexual stage of their complex life cycle, which determines transmission in the mosquito vector (Morrissette and Sibley, 2002; Portman and Slapeta, 2014). Flagella formation in *Plasmodium spp* is unique compared with that of other eukaryotes, including the closely related Apicomplexa, as they arise from *de novo* formed basal bodies and are directly assembled in the male gametocyte cytoplasm, and thus do not require building material to be transported by the IFT mechanisms of many other systems (Briggs et al., 2004; Sinden et al., 1976; Sinden et al., 2010). This assembly mechanism is rapid and presumably an adaptation to facilitate efficient transition through this life cycle stage.

We show that *Plasmodium* kinesin-8B has a specific role in building the male gamete flagellum, and in its absence parasite transmission from insect vector to mammalian host is blocked. The severe defect in exflagellation exhibited by *Δkinesin-8B* parasites is consistent with the location of protein we describe, and with its presence in the male proteome (Khan et al., 2005; Talman et al., 2014). Furthermore, a specific role for kinesin-8B at this life cycle stage is consistent with it being dispensable during blood stages (this study and (Bushell et al., 2017)).

Male gamete development in the malaria parasite involves three rounds of replication and mitotic division followed by chromosome condensation and cytokinesis, resulting in eight gametes (Sinden et al., 2010; Guttery et al., 2012a). DNA content analysis of the *Δkinesin-8B* mutant showed that the normal three rounds of replication occurred since octaploid nuclei (8N) were observed at 8-10 mpa. The distinct separation of kinesin-8B from the kinetochore marker Ndc80, and the normal nuclear ultrastructure, further substantiate the idea that this kinesin is not part of the mitotic spindle within the nucleus. The experiments using gametocyte-specific inhibitors (Delves et al., 2018; Gamo et al., 2010), provide further discrimination between MT-dependent events in the nucleus – where these compounds have no effect on Ndc80 localisation and therefore presumably no effect on mitotic spindle MTs – and the cytoplasm, where they have considerable effect on cytoplasmic axoneme assembly and kinesin-8B localisation.

Global transcript analysis and qRT-PCR examination of genes coding for proteins shown to affect cytokinesis and chromosome condensation, like CDPK4, CDC20, GEST and MAP2 (Billker et al., 2004; Guttery et al., 2012; Talman et al., 2011; Tewari et al., 2005) revealed no differentially expressed genes in the mutant, in defective *Δkinesin-8B* parasite male gametogenesis which were deemed important. This suggests that the observed phenotype is due to the absence of kinesin-8B alone.

The ultrastructure studies provide a very striking comparison between the assembled axonemes of wild type parasites and the lack of proper axoneme assembly in the absence of kinesin-8B. Examination of two different time points (8 and 15 mpa) revealed that single and doublet MTs were formed and elongated in the mutant parasites, but they were not organised into the classical 9+2 symmetry. Furthermore, although the amorphous nature of the basal body precluded identification of structural abnormalities in the absence of kinesin-8B, there appeared to be at least partial loss of the tight association between the basal body and the nuclear pole. This phenotype is very different to that of many other mutants with exflagellation defects (Billker et al., 2004; Ferguson DJP, 2013; Wall et al., 2018), but there is some similarity to that of other mutants affecting the basal body (SAS6) or involved in assembly of the central pair of MTs (PF16) in *Plasmodium* axonemes (Marques et al., 2015; Straschil et al., 2010). These data suggest that the kinesin-8B deletion causes a defect in both the basal body and axonemes.

Using live cell imaging, we show the dynamic localisation of kinesin-8B from the earliest stages of male gamete development (summarized in Fig 5). Kinesin-8B is a cytoplasmic protein exhibiting a basal body-like pattern in the early stages of gamete development – this highlights that basal body replication occurs within 2 minutes of activation and that associated axoneme-like structures are assembled later. The association of kinesin-8B with axonemes is supported by co-localisation with cytoplasmic MT. The pattern of axoneme development revealed by tracking kinesin-8B is consistent with earlier ultrastructural studies (Sinden et al 1976, 1978). The formation of basal bodies from a single MTOC-like structure was evident at a very early time in gamete development (Sinden et al., 1978; Sinden et al., 1976). Here we have displayed the real-time dynamics of their duplication and localization as distinct structures. These features are consistent with the basal body defects resulting in disorganised axoneme assembly that we observe with the deletion of kinesin8B.

Kinesin motors participate in diverse functions in many processes including cell division, intracellular transport, intraflagellar transport and controlling MT dynamics ((Cross and McAinsh, 2014; Dawson et al., 2007; Scholey, 2008; Verhey and Hammond, 2009). The lack of association of kinesin-8B with mitotic assembly within the nucleus - and with kinetochores in particular - is distinct from what is seen for kinesin-8As in most eukaryotes (Savoian et al., 2004; Savoian and Glover, 2010; Wang et al., 2016). The metazoan kinesin-8B, Kif19, is involved in ciliary length regulation, but the role of kinesin-8B in axonemal assembly appears to be more profound in *Plasmodium* than controlling length alone. Kinesin-8B is consistently present in those Apicomplexa with flagellated gametes like *Plasmodium*, *Toxoplasma* and *Eimeria* and absent in other genera like *Theileria, Babesia* and *Cryptosporidium* which lack flagellated gametes (Zeeshan M, 2019). The properties of kinesin-8B contrast with those of *Plasmodium* kinesin-8X, which is located on the mitotic spindle (Zeeshan M, 2019).

In conclusion, this is the first study exploring the real-time dynamics and functional role of the kinesin-8B molecular motor in flagellum formation during *Plasmodium* male gamete development. It has a key role in establishing basal body formation and axoneme structure and assembly, thereby regulating male gamete development, which is an essential stage in parasite transmission.

## Materials and Methods

### Ethics statement

The animal work performed in the UK passed an ethical review process and was approved by the United Kingdom Home Office. Work was carried out under UK Home Office Project Licenses (40/3344 and 30/3248) in accordance with the United Kingdom ‘Animals (Scientific Procedures) Act 1986’. Six to eight week-old female Tuck-Ordinary (TO) (Harlan) outbred mice were used for all experiments in the UK.

### Generation of transgenic parasites

The gene-deletion targeting vector for *kinesin-8B* (PBANKA_020270) *was* constructed using the pBS-DHFR plasmid, which contains polylinker sites flanking a *T. gondii dhfr/ts* expression cassette conferring resistance to pyrimethamine, as described previously (Saini et al., 2017). PCR primers N1261 and N1262 were used to generate a 1000 bp fragment of *kinesin-8B* 5′ upstream sequence from genomic DNA, which was inserted into *Apa*I and *Hin*dIII restriction sites upstream of the *dhfr/ts* cassette of pBS-DHFR. A 1240 bp fragment generated with primers N1263 and N1264 from the 3′ flanking region of *kinesin-8B* was then inserted downstream of the *dhfr/ts* cassette using *Eco*RI and *Xba*I restriction sites. The linear targeting sequence was released using *Apa*I/*Xba*I. A schematic representation of the endogenous *Pbkinesin-8B* locus the constructs and the recombined *kinesin-8B* locus can be found in S1 Fig.

The C-terminus of kinesin-8B was tagged with GFP by single crossover homologous recombination in the parasite. To generate the kinesin-8B-GFP line, a region of the *kinesin-8* gene downstream of the ATG start codon was amplified using primers T1991 and T1992, ligated to p277 vector, and transfected as described previously (Guttery et al, 2012). A schematic representation of the endogenous *kinesin-8B* locus (PBANKA_020270), the constructs and the recombined *kinesin-8B* locus can be found in S2 Fig. The oligonucleotides used to generate the mutant parasite lines can be found in S2 Table. *P. berghei* ANKA line 2.34 (for GFP-tagging) or ANKA line 507cl1 expressing GFP (for gene deletion) were transfected by electroporation (Janse et al., 2006).

### Parasite genotype analyses

For the gene knockout parasites, diagnostic PCR was used with primer 1 (IntN126) and primer 2 (ol248) to confirm integration of the targeting construct, and primer 3 (N102 KO1) and primer 4 (N102 KO2) were used to confirm deletion of the *kinesin-8B* gene (S1 Fig). For the parasites expressing a C-terminal GFP-tagged kinesin-8B protein, diagnostic PCR was used with primer 1 (IntT199) and primer 2 (ol492) to confirm integration of the GFP targeting construct (S2 Fig).

### Purification of gametocytes

The purification of gametocytes was achieved using a protocol described previously (Beetsma et al., 1998) with some modifications. Briefly, parasites were injected into phenylhydrazine treated mice and enriched by sulfadiazine treatment after 2 days of infection. The blood was collected on day 4 after infection and gametocyte-infected cells were purified on a 48% v/v NycoDenz (in PBS) gradient. (NycoDenz stock solution: 27.6% w/v NycoDenz in 5 mM Tris-HCl, pH 7.20, 3 mM KCl, 0.3 mM EDTA). The gametocytes were harvested from the interface and washed.

### Live cell- and -time lapse imaging

Purified gametocytes were examined for GFP expression and localization at different time points (0, 1-15 min) after activation in ookinete medium (Guttery et al 2014). Images were captured using a 63x oil immersion objective on a Zeiss Axio Imager M2 microscope fitted with an AxioCam ICc1 digital camera (Carl Zeiss, Inc). Time-lapse videos (1 frame every 5 sec for 15-20 cycles) were taken with a 63X objective lens on the same microscope and analysed with the AxioVision 4.8.2 software.

### Fixed immunofluorescence assay and measurements

The purified gametocytes from kinesin-8B-GFP, WT-GFP and *Δkinesin-8B* parasites were activated in ookinete medium then fixed at 0 min, 1-2 min, 6-8 min and 15 min post-activation with 4% paraformaldehyde (PFA, Sigma) diluted in microtubule stabilising buffer (MTSB) for 10-15 min and added to poly-L-lysine coated slides. Immunocytochemistry was performed using primary GFP-specific rabbit monoclonal antibody (mAb) (Invitrogen-A1122; used at 1:250) and primary mouse anti-α tubulin mAb (Sigma-T9026; used at 1:1000). Secondary antibodies were Alexa 488 conjugated anti-mouse IgG (Invitrogen-A11004) and Alexa 568 conjugated anti-rabbit IgG (Invitrogen-A11034) (used at 1 in 1000). The slides were then mounted in Vectashield 19 with DAPI (Vector Labs) for fluorescence microscopy. Parasites were visualised on a Zeiss AxioImager M2 microscope fitted with an AxioCam ICc1 digital camera (Carl Zeiss, Inc).

To measure nuclear DNA content of activated microgametocytes by direct immunofluorescence, images of parasites fixed (0 mpa and 8-10 mpa) and stained as above were analyzed using the ImageJ software (version 1.44) (National Institute of Health) as previously described (Tewari et al., 2005).

### Generation of dual tagged parasite lines

The kinesin-8B-GFP parasites were mixed with Ndc80-cherry parasites in equal numbers and injected into a mouse. Mosquitoes were fed on this mouse 4 to 5 days after infection when gametocyte parasitemia was high. These mosquitoes were checked for oocyst development and sporozoite formation at day 14 and day 21 after feeding. Infected mosquitoes were then allowed to feed on naïve mice and after 4 - 5 days these mice were examined for blood stage parasitemia by microscopy with Giemsa-stained blood smears. In this way, some parasites expressed both kinesin-8B-GFP and Ndc80-cherry in the resultant gametocytes, and these were purified and fluorescence microscopy images were collected as described above.

### Inhibitor studies

Gametocytes were purified as above and treated with two antimalarial molecules (TCMDC-123880 and DDD01028076) (Delves et al., 2018; Gamo et al., 2010) at 4 mpa and then fixed with 4% PFA at 8 min after activation. Dimethyl sulfoxide (DMSO) was used as a control treatment. These fixed gametocytes were then examined on a Zeiss AxioImager M2 microscope fitted with an AxioCam ICc1 digital camera (Carl Zeiss, Inc).

### Parasite phenotype analyses

Blood containing approximately 50,000 parasites of the *Δkinesin-8B* line was injected intraperitoneally (i.p) into mice to initiate infections. Asexual stages and gametocyte production were monitored by microscopy on Giemsa stained thin smears. Four to five days post infection, exflagellation and ookinete conversion were examined as described previously (Guttery et al, 2012) with a Zeiss AxioImager M2 microscope (Carl Zeiss, Inc) fitted with an AxioCam ICc1 digital camera. To analyse mosquito transmission, 30–50 *Anopheles stephensi* SD 500 mosquitoes were allowed to feed for 20 min on anaesthetized, infected mice with an asexual parasitemia of 15% and a comparable numbers of gametocytes as determined on Giemsa stained blood films. To assess mid-gut infection, approximately 15 guts were dissected from mosquitoes on day 14 post feeding, and oocysts were counted on an AxioCam ICc1 digital camera fitted to a Zeiss AxioImager M2 microscope using a 63x oil immersion objective. On day 21 post-feeding, another 20 mosquitoes were dissected, and their guts crushed in a loosely fitting homogenizer to release sporozoites, which were then quantified using a haemocytometer or used for imaging. Mosquito bite back experiments were performed 21 days post-feeding using naive mice, and blood smears were examined after 3-4 days.

### Electron microscopy

Gametocytes activated for 8 min and 15 min were fixed in 4% glutaraldehyde in 0.1 M phosphate buffer and processed for electron microscopy as previously described (Ferguson et al., 2005). Briefly, samples were post fixed in osmium tetroxide, treated *en bloc* with uranyl acetate, dehydrated and embedded in Spurr’s epoxy resin. Thin sections were stained with uranyl acetate and lead citrate prior to examination in a JEOL1200EX electron microscope (Jeol UK Ltd).

### Quantitative Real Time PCR (qRT-PCR) analyses

RNA was isolated from gametocytes using an RNA purification kit (Stratagene). cDNA was synthesised using an RNA-to-cDNA kit (Applied Biosystems). Gene expression was quantified from 80 ng of total RNA using a SYBR green fast master mix kit (Applied Biosystems). All the primers were designed using the primer3 software (Primer-blast, NCBI). Analysis was conducted using an Applied Biosystems 7500 fast machine with the following cycling conditions: 95°C for 20 s followed by 40 cycles of 95°C for 3 s; 60°C for 30 s. Three technical replicates and three biological replicates were performed for each assayed gene. The *hsp70* (PBANKA_081890) and *arginyl-t RNA synthetase* (PBANKA_143420) genes were used as endogenous control reference genes. The primers used for qPCR can be found in **S2 Table**.

### RNAseq analysis

Libraries were prepared from lyophilized total RNA using the KAPA Library Preparation Kit (KAPA Biosystems) following amplification for a total of 12 PCR cycles (12 cycles [15 s at 98°C, 30 s at 55°C, 30 s at 62°C]) using the KAPA HiFi HotStart Ready Mix (KAPA Biosystems). Libraries were sequenced using a NextSeq500 DNA sequencer (Illumina), producing paired-end 75-bp reads.

FastQC (https://www.bioinformatics.babraham.ac.uk/projects/fastqc/), was used to analyze raw read quality, and based on this information, the first 10 bp of each read and any adapter sequences were removed using Trimmomatic (http://www.usadellab.org/cms/?page=trimmomatic). The resulting reads were mapped against the *P. berghei* ANKA genome (v36) using HISAT2 (version 2-2.1.0), using default parameters. Uniquely mapped, properly paired reads were retained using SAMtools (http://samtools.sourceforge.net/). Genome browser tracks were generated and viewed using the Integrative Genomic Viewer (IGV) (Broad Institute). Raw read counts were determined for each gene in the *P. berghei* genome using BedTools (https://bedtools.readthedocs.io/en/latest/#) to intersect the aligned reads with the genome annotation. Differential expression analysis was done using DESeq2 to calculate log_2_fold expression changes between the knockout and wild-type conditions for every gene. Gene ontology enrichment was done either using R package TopGO, or on PlasmoDB (http://plasmodb.org/plasmo/) with repetitive terms removed by REVIGO (http://revigo.irb.hr/).

### Statistical analysis

All statistical analyses were performed using GraphPad Prism 7 (GraphPad Software). For qRT-PCR, an unpaired t-test was used to examine significant differences between wild-type and mutant strains.

## Data Availability

Sequence reads have been deposited in the NCBI Sequence Read Archive with accession number: PRJNA549466

## Acknowledgments.

We thank Julie Rodgers for helping to maintain the insectary and other technical works.

## Funding

This work was supported by: MRC UK (G0900278, MR/K011782/1) and BBSRC (BB/N017609/1) to RT and MZ; the BBSRC (BB/N018176/1) to CAM; the Francis Crick Institute (FC001097), the Cancer Research UK (FC001097), the UK Medical Research Council (FC001097), and the Wellcome Trust (FC001097) to AAH; Wellcome Trust Equipment Grant to DJPF; the NIH/NIAID (R01 AI136511) and the University of California, Riverside (NIFA-Hatch-225935) to KGLR.

## Author Contributions

Conceived and designed the experiments: RT and CAM. Performed the Functional and GFP tagging experiments: RT MZ ER ED DB. RNAseq experiment: KGLR SA TM. QRT-PCR analysis: MZ DB. Transmission electron microscopy: DAJF AB SV. Analyzed the data: RT MZ SV DAJF SA KGLR CAM AAH. Contributed reagents/materials/analysis tools: RT DAJF KGLR SV MD. Writing - Original Draft, MZ and RT; Writing - Review & Editing, CAM AAH KGLR DAJF SA, MZ, RT and all others contributed.

## Supplementary materials

### Supplementary Figures

**S1 Fig:**
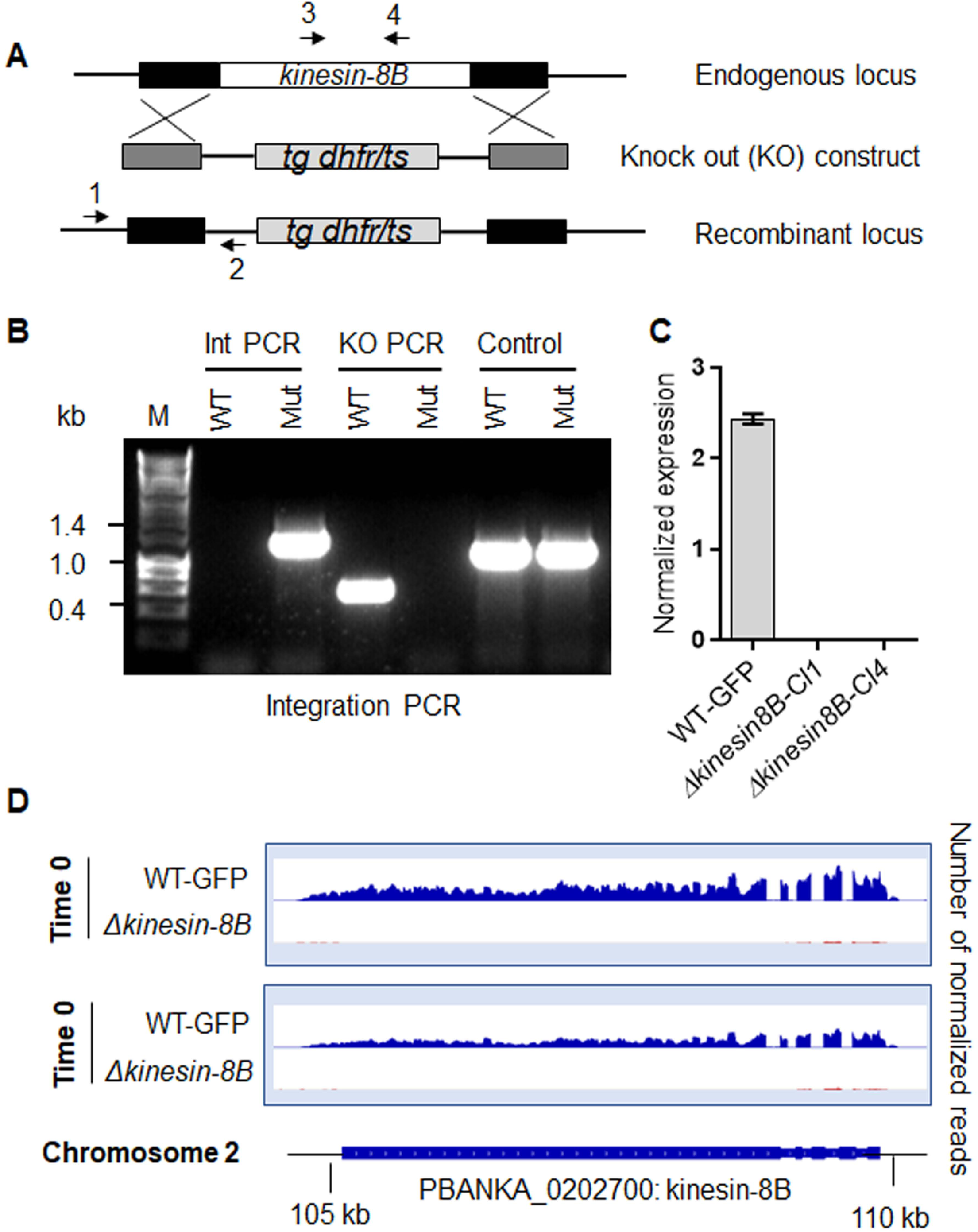
Generation and genotypic analysis of *Δkinesin-8B* parasites. **A**. Schematic representation of the endogenous kinesin-8B locus, the targeting knockout construct and the recombined kinesin-8B locus following double homologous crossover recombination. Arrows 1 and 2 indicate PCR primers used to confirm successful integration in the kinesin-8B locus following recombination and arrows 3 and 4 indicate PCR primers used to show deletion of the kinesin-8B gene. **B.** Integration PCR of the kinesin-8B locus in WT-GFP and *Δkinesin-8B* (Mut) parasites using primers INT N105 and ol248. Integration of the targeting construct gives a band of 1.5 kb. **C.** qRT-PCR analysis of transcript in WT-GFP and *Δkinesin-8B* parasites. **D.** RNAseq analysis confirming lack of transcripts from locus on chromosome 2 in *Δkinesin-8B* parasites.

**S2 Fig:**
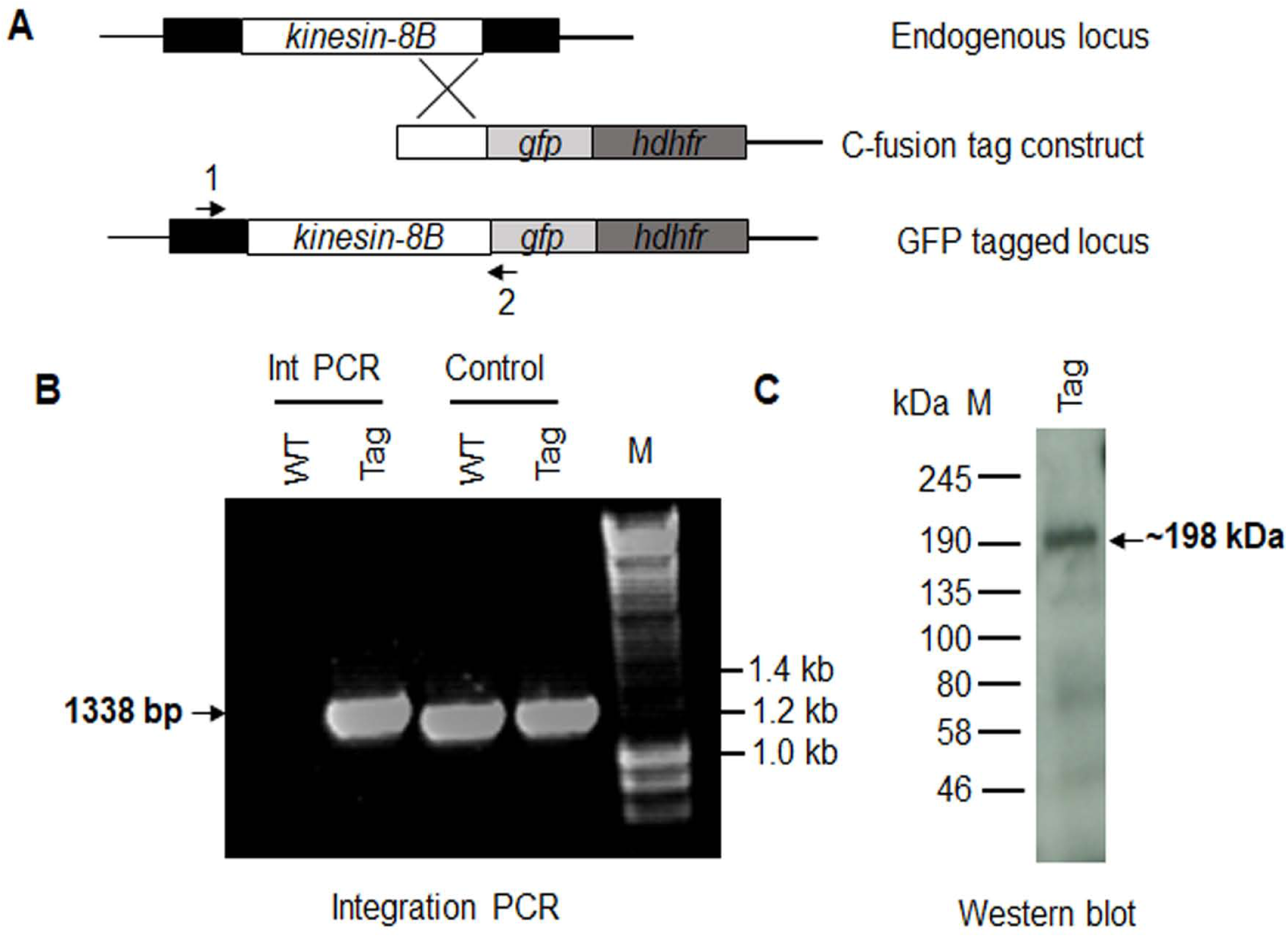
Generation and genotypic analysis of kinesin-8B-GFP. **A**. Schematic representation of the endogenous *pbkinesin-8B* locus, the GFP-tagging construct and the recombined *kinesin-8B* locus following single homologous recombination. Arrows 1 and 2 indicate the position of PCR primers used to confirm successful integration of the construct. **B**. Diagnostic PCR of *kinesin-8B* and WT parasites using primers IntT199 (Arrow 1) and ol492 (Arrow 2). Integration of the kinesin-8B tagging construct gives a band of 1338 bp. Tag = kinesin-8B-GFP parasite line. **C**. Western blot of kinesin-8B-GFP (∼198 kDa) protein illustrates the presence of the protein in gametocyte stage.

### Supplementary Tables

<u>S1 Table.</u> List of differentially expressed genes between Δ*kinesin-8B* and WT activated gametocytes

<u>S2 Table.</u> Oligonucleotides used in this study

### Supplementary Videos

<u>SV1 Video.</u> Time lapse video showing duplicated tetrad foci of kinesin-8B moving apart from each other in 50 sec.

<u>SV1 Video.</u> Dynamic movement of duplicated tetrad foci and emerging fibre-like structure of kinesin-8B in 50 sec.

## Notes

#### Summary of Updates

Few changes in text, Figure 2 legend and references

